# Comparative analysis of ionic strength tolerance between freshwater and marine Caulobacterales adhesins

**DOI:** 10.1101/523142

**Authors:** Nelson K. Chepkwony, Cécile Berne, Yves V. Brun

## Abstract

Bacterial adhesion is affected by environmental factors, such as ionic strength, pH, temperature, and shear forces, and therefore marine bacteria must have developed holdfasts with different composition and structures than their freshwater counterparts to adapt to their natural environment. The dimorphic *α*-proteobacterium *Hirschia baltica* is a marine budding bacterium in the Caulobacterales clade. *H*. *baltica* uses a polar adhesin, the holdfast, located at the cell pole opposite the reproductive stalk for surface attachment and cell-cell adhesion. The holdfast adhesin has been best characterized in *Caulobacter crescentus*, a freshwater member of the Caulobacterales, and little is known about holdfast composition and properties in marine Caulobacterales. Here we use *H. baltica* as a model to characterize holdfast properties in marine Caulobacterales. We show that freshwater and marine Caulobacterales use similar genes in holdfast biogenesis and that these genes are highly conserved among the two genera. We also determine that *H. baltica* produces larger holdfast than *C. crescentus* and that those holdfasts have a different chemical composition, as they contain N-acetylglucosamine and galactose monosaccharide residues and proteins, but lack DNA. Finally, we show that *H. baltica* holdfasts tolerate higher ionic strength than those of *C. crescentus*. We conclude that marine Caulobacterales holdfasts have physicochemical properties that maximize binding in high ionic strength environments.

**IMPORTANCE:** Most bacteria spend a large amount of their lifespan attached to surfaces, forming complex multicellular communities called biofilms. Bacteria can colonize virtually any surface, therefore they have adapted to bind efficiently in very different environments. In this study, we compare the adhesive holdfasts produced by the freshwater bacterium *C. crescentus* and a relative, the marine bacterium *H. baltica*. We show that *H. baltica* holdfasts have a different morphology and chemical composition, and tolerate high ionic strength. Our results show that *H. baltica* holdfast is an excellent model to study the effect of ionic strength on adhesion and providing insights on the physicochemical properties required for adhesion in the marine environment.

## INTRODUCTION

In their natural environments, bacteria preferentially form surface-associated communities, known as biofilms (1). To irreversibly adhere to surfaces and form these complex mutil-cellular communities, bacteria produce strong adhesins, mainly composed of proteins or polysaccharides (2, 3). Bacterial adhesion is affected by different environmental conditions such as pH, temperature, shear forces, and ionic strength (2, 4-6). In marine environments, bacteria face 500 times higher ionic strength than in freshwater (7), therefore, marine bacteria have evolved ways to overcome the effect of ionic strength and bind permanently to surfaces in high salt environments such as seas and oceans.

Caulobacterales are Alphaproteobacteria found in various habitats, from oligotrophic aquatic and nutrient-rich soil environments (8, 9). The aquatic Caulobacterales species live in a wide range of environments with different salinity levels, such as pristine fresh river and lake waters, brackish ponds, and marine waters, making them a good model for studying bacterial adhesion in different ionic environments. Caulobacterales species use a polar adhesin structure called the holdfast to adhere permanently to surfaces and form biofilms (8, 10, 11). The holdfast has been primarily studied in *Caulobacter crescentus*, a freshwater member of the Caulobacterales (2, 3, 12). The *C. crescentus* holdfast uses both electrostatic and hydrophobic interactions to attach to different surfaces (6). The binding affinity of the *C. crescentus* holdfast is dramatically impaired in the presence of NaCl (6), yet marine Caulobacterales adhere to surfaces at considerably higher ionic strength, suggesting that their holdfasts have different properties. However, little is known about holdfasts from marine Caulobacterales and the molecular mechanism used to adhere successfully to surfaces in salinity environments is currently unknown.

*C. crescentus* holdfast is the strongest characterized bioadhesive, with an adhesion force of 70N / mm^2^ (13). Despite being first identified almost 85 years ago (14), the exact composition and structure of the *C. crescentus* holdfast remains elusive. Wheat Germ Agglutinin (WGA) lectin-binding assays show that the holdfast contains N-acetylglucosamine (GlcNAc) residues (10), while other studies suggest that the holdfast is also composed of unidentified peptide and DNA residues (15). The *C. crescentus* holdfast polysaccharide is produced via a polysaccharide synthesis and export pathway similar to the group I capsular polysaccharide synthesis Wzy/Wzx-dependent pathway in *E. coli* (16, 17) (Fig. 1A). Holdfast polysaccharide synthesis is thought to be initiated in the cytoplasm by a glycosyltransferase HfsE, which transfers activated sugar phosphate from uridine diphosphate (UDP-GlcNAc) to undecaprenyl-phosphate (Und-P) lipid carrier. Additional sugar residues are then added to form a repeat unit on the lipid carrier by three glycosyltransferases HfsG, HfsJ (17) and HfsL (18). The acetyltransferase HfsK (32) and the polysaccharide deacetylase HfsH (21) modify one or more sugar residue. The lipid carrier with the repeat units is transported across the inner membrane into the periplasm by a flippase (HfsF) (17, 19). In the periplasm, the repeat units are polymerized by two polysaccharide polymerases HfsC and HfsI (17). The holdfast polysaccharide chain is then secreted through the export proteins complex, composed of HfsA, HfsB and HfsD (20-22). Once outside the cell, holdfast polysaccharides are anchored unto the cell envelop by the action of <u>h</u>oldfast <u>a</u>nchor (*hfa*) proteins HfaA, HfaB, HfaD and HfaE (18, 23-25).

**Figure 1:**
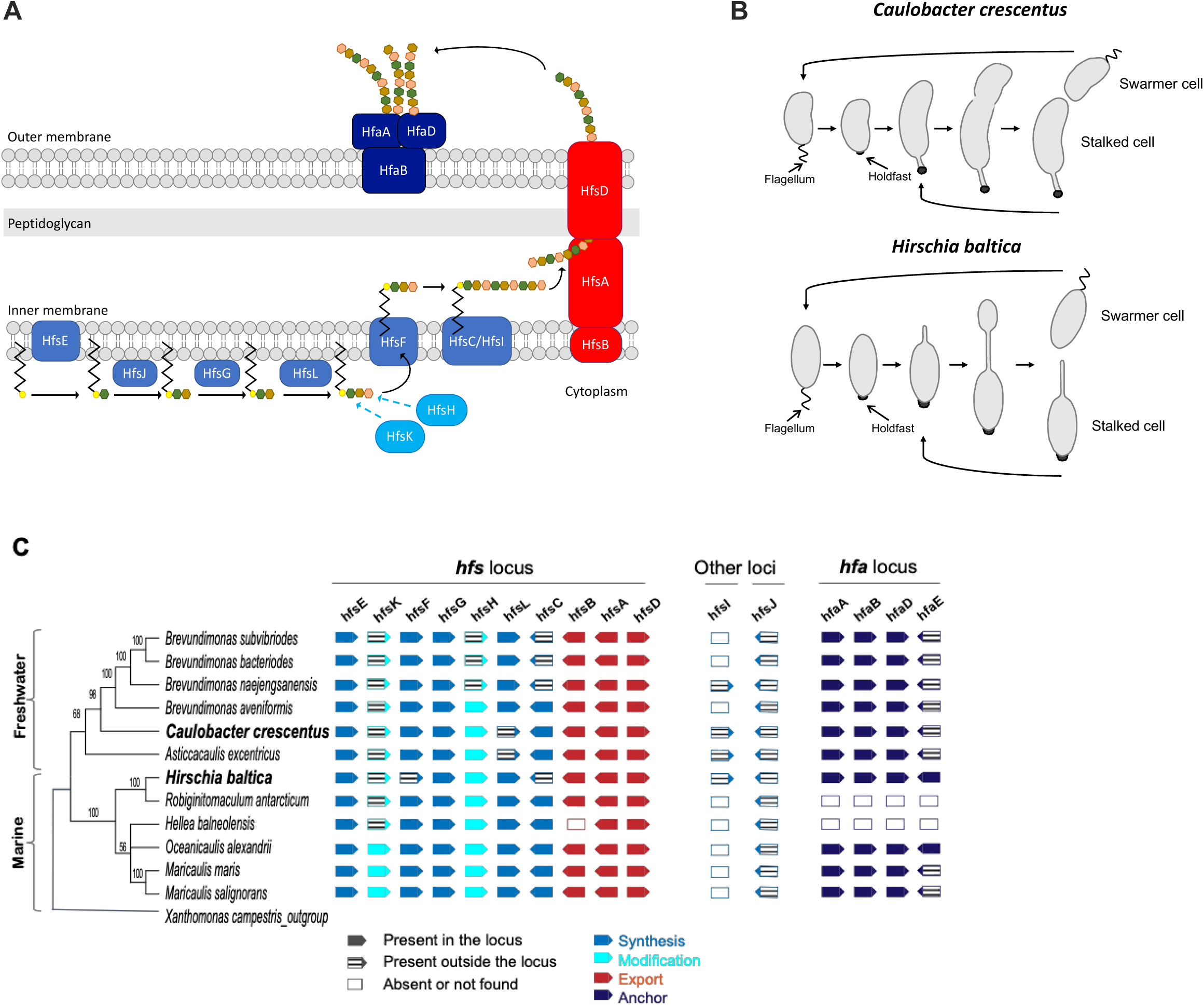
Organization of the holdfast gene cluster in *H. baltica*. **A.** Schematic of holdfast synthesis, modification, secretion, and anchor machineries. Holdfast polysaccharide synthesis is initiated by glycosyltransferase HfsE, which transfers activated sugar precursors in the cytoplasm to a lipid carrier. Three glycosyltransferases HfsJ, HfsG and HfsL add different sugars to the growing polysaccharide. The acetyltransferase HfsK and the deacetylase HfsH modify one or more sugar residue and then a flippase HfsF transports the lipid carrier into periplasm. Repeat units are polymerized by polymerases HfsC and HfsI. The polysaccharide is exported to the outside of the cell through the HfsA, HfsB, HfsD complex. The exported polysaccharide is then anchored to the cell body by secreted protein HfaA, HfaB and HfaD. The different colors of hexagons represent different sugars. **B.** Diagrams of *C. crescentus* and *H. baltica* dimorphic cell cycles. A motile swarmer cell differentiates into stalked cell by shedding its flagellum and synthesizing holdfast at the same cell pole. *C. crescentus* stalked cell divide asymmetrical to produce a motile swarmer and a stalked cell (top panel) and *H. baltica* reproduces by budding a motile swarmer off the stalk (bottom panel). **C.** Maximum likelihood phylogeny inferred from 16S rRNA sequences of selected freshwater and marine members of Caulobacterales. Node values represent clade frequency of 1000 bootstraps. The genes were identified using reciprocal best hit analysis on fully sequenced Caulobacterales genomes. Solid gene symbols indicate genes within the *hfs* or *hfa* loci while striped symbols indicate the genes translocated from these loci to a different location in the genome. Empty boxes indicate absent or missing genes. Blue represents synthesis genes, turquoise represents modification genes, red represents polysaccharide export genes and navy blue represents anchor genes.

*Hirschia baltica* is a marine Caulobacterale isolated from surface water taken from a boat landing in the Kiel Fjord inlet of the Baltic sea (Germany) (26). *H. baltica* has a dimorphic life-cycle similarly to *C. crescentus* (26), but reproduces by budding from the tip of the stalk (Fig.1B). Newborn swarmer cells are motile by means of a polar flagellum, and differentiate into sessile stalked cells after flagellum ejection. The sessile cells produce a holdfast at the same pole as the flagellum and synthesize a stalk at the opposite pole (27). *H. baltica* have been shown to produce holdfast containing GlcNAc residues, using fluorescent WGA lectin (27, 28). The vast majority of studies on holdfast have been done using *C. crescentus,* therefore *H. baltica* holdfast is poorly understood.

As bacteria have to develop different strategies to adhere to surfaces in a given environment, we hypothesized that *H. baltica* produces holdfasts with different physicochemical properties because *H. baltica* natural habitat is high ionic strength sea water (26), while the freshwater *C. crescentus* holdfast is highly sensitive to salt (6). Here we study *H. baltica* holdfast composition and properties. Using both genetics and bioinformatics analyses, we show that freshwater and marine Caulobacterales use orthologous genes in holdfast biogenesis and these genes are highly conserved among these genera. We show that *H. baltica* produces more holdfast material than *C. crescentus* and that the holdfasts of the two genera have a different chemical composition and behave differently. In addition to GlcNAc monosaccharides, we show that *H. baltica* holdfasts also contain galactose residues, and uncharacterized peptides different than the ones found in *C. crescentus* holdfasts. Finally, we demonstrate that *H. baltica* holdfast tolerates higher ionic strength than that of *C. crescentus*.

## RESULTS

### Organization of the holdfast genes in *H. baltica*

The genes essential for holdfast synthesis and export in the *C. crescentus hfs* locus (*hfsG, hfsB, hfsA* and *hfsD*) are conserved in *H. baltica* (27, 28). To determine if the genomic organization of all the known holdfast related genes is conserved in both species, we performed reciprocal best hit analyses using the *C. crescentus hfs* (holdfast synthesis, modification, and export) and *hfa* (holdfast anchoring) genes (Fig. 1C). We also extended our analysis to other fully sequenced available Caulobacterales genomes for a more global overview of the organization of these genes in this clade (Fig. 1C). Table 1 gives the locus tag names of all the holdfast related genes used in this study for *C. crescentus* CB15 (29), *C. crescentus* NA1000 (30), and *H. baltica* IFAM 1418^T^ (27) type strains.

**Table 1:** Genes involved in holdfast synthesis, modification and anchoring

| Gene name | *C. crescentus*<br>CB15 | *C. crescentus*<br>NA1000 | *H. baltica* strain<br>IFAM 1418<sup>T</sup> |
| --- | --- | --- | --- |
| **Export apparatus** |  |  |  |
| *hfsA* | CC2431 | CCNA_02513 | Hbal_1968 |
| *hfsB* | CC2430 | CCNA_02512 | Hbal_1967 |
| *hfsD* | CC2432 | CCNA_02514 | Hbal_1969 |
| **Synthesis genes** |  |  |  |
| *hfsC* | CC2429 | CCNA_02511 | Hbal_1972 |
| *hfsE* | CC2425 | CCNA_02507 | Hbal_1963 |
| *hfsJ* | CC0095 | CCNA_00094 | Hbal_1784 |
| *hfsG* | CC2427 | CCNA_02509 | Hbal_1964 |
| *hfsL* | CC2277 | CCNA_02361 | Hbal_1966 |
| *hfsI* | CC0500 | CCNA_00533 | Hbal_2115 |
| *hfsF* | CC2426 | CCNA_02508 | Hbal_0100 |
| **Modification genes (non essential)** |  |  |  |
| *hfsH* | CC2428 | CCNA_02510 | Hbal_1965 |
| *hfsK* | CC3689 | CCNA_03803 | Hbal_0069 |
| **Anchor genes (non essential)** |  |  |  |
| *hfaA* | CC2628 | CCNA_02711 | Hbal_0652 |
| *hfaB* | CC2630 | CCNA_02712 | Hbal_0651 |
| *hfaD* | CC2629 | CCNA_02713 | Hbal_0650 |
| *hfaE* | CC2639 | CCNA_02722 | Hbal_0649 |

All the genes reported involved in holdfast synthesis in *C. crescentus* are present in the analyzed genomes, with a few rearrangements (Fig. 1C). The general organization of the *hfs* locus is conserved in all the Caulobacterales genomes analyzed, with the genes encoding proteins essential for holdfast synthesis (glycosyltransferase gene *hfsG* and export genes *hfsA, hfsB*, and *hfsD*) and the initiating glycosyltransferase gene *hfsE* in a similar organization as in *C. crescentus*. Some of the genes involved in holdfast synthesis and modification in *C. crescentus* are not part of the *hfs* gene cluster (genes encoding the polymerase HfsI (17), the glycotransferases HfsJ (31) and HfsL (18), and N-acetyltransferase HfsK (32)); these genes are also present in *H. baltica.* Interestingly, in the genomes of the marine Caulobacterales *Oceanicaulis alexandrii, Maricaulis maris and Maricaulis salignorans,* all the *hfs* genes are found in one locus except *hfsJ* (Fig. 1C). This suggests that the ancestral *hfs* locus might have contained most of the *hfs* genes. Most of the genomes analyzed had only one polysaccharide polymerase gene, *hfsC* while other have a paralogous polysaccharide polymerase gene *hfsI* (Fig. 1C) (17).

Once exported outside the cell by the HfsDAB complex, the holdfast is anchored to the cell envelope by the action of anchor proteins that have been identified and characterized in *C. crescentus* HfaA, HfaB, and HfaD (18, 23-25). The organization of the three anchor genes *hfaA, hfaB* and *hfaD* in the *hfa* is conserved in all the analyzed Caulobacterales genomes (Fig. 1C). In *C. crescentus* and most of the tested Caulobacterales, the recently identified holdfast anchor gene *hfaE* (18) is not part of the *hfaABD* operon, while it is present in the *hfa* locus in both *H. baltica* and *Oceanicaulis alexandrii* (Fig. 1C). We could not find orthologs of the *hfa* genes in the genomes of *Robiginitomaculum antarticum* and *Hellea balneolensis*, but this may be due to the incomplete nature of the genome. Alternatively these species have a different mechanism to anchor holdfast to the surface of the cell, as is the case for several other Alphaproteobacteria (33, 34).

### Role of the *hfs and hfa* genes in *H. baltica*

To determine if the genes identified in Fig. 1C are involved in holdfast production and anchoring in *H. baltica*, we created in-frame deletion mutants of the *hfa* genes encoding the anchor proteins, and the *hfs* genes shown to be essential for holdfast synthesis in *C. crescentus* (12). We first monitored the presence of holdfasts in these mutants using fluorescence microscopy with fluorescently labelled WGA lectin (10) (Fig. 2A). We also quantified biofilm formation after 12 h of incubation at room temperature on a plastic surface, using 24-well PVC plates (Fig. 2B). All mutants could be complemented *in trans* by a replicating plasmid encoding a copy of the deleted gene (Fig. 2B).

**Figure 2:**
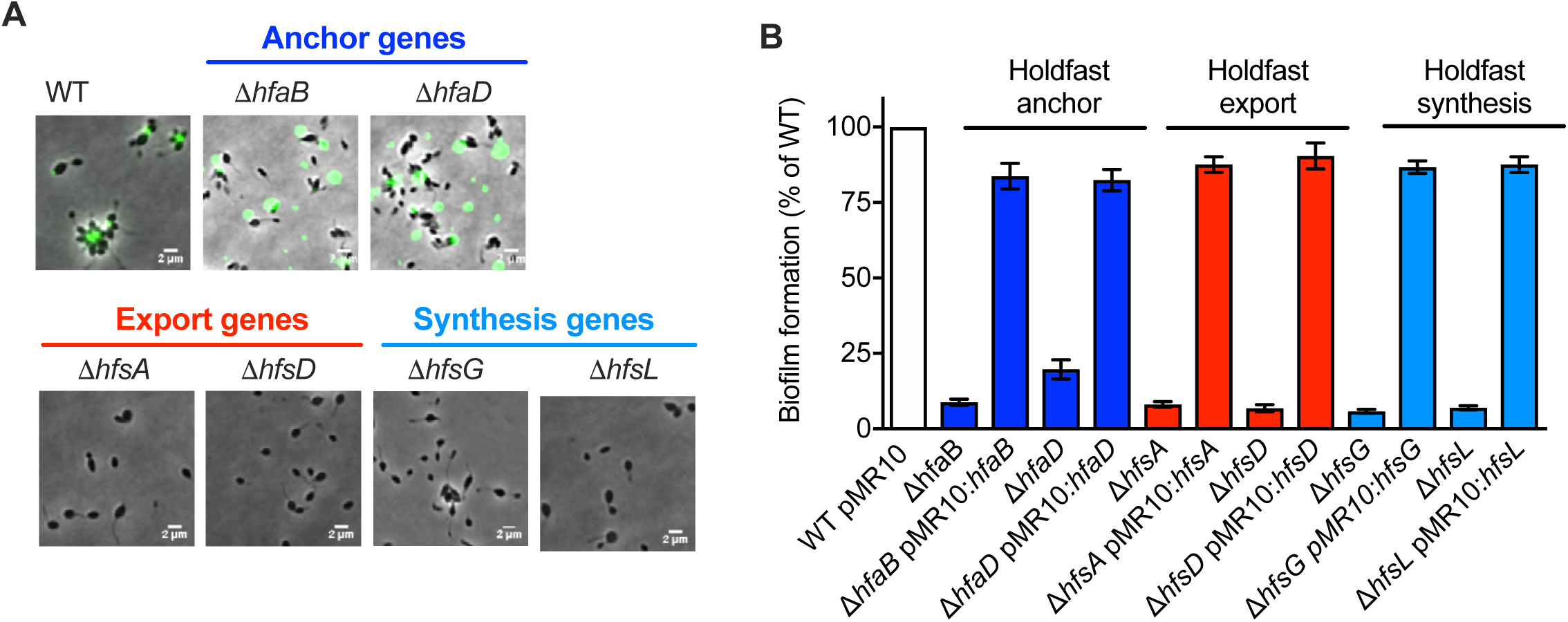
Role of the *hfs* and *hfa* genes in *H. baltica* holdfast production. **A.** Representative images showing merged phase and fluorescence channels of different *H. baltica* WT and mutant strains with holdfast labeled with WGA-AF488 (green): *H. baltica* holdfast anchor mutants Δ*hfaB* and *ΔhfaD,* export mutants Δ*hfsA* and Δ*hfsD,* and synthesis mutants Δ*hfsG* and *ΔhfsL.* **B.** Quantification of biofilm using crystal violet assay after 12 hours for *hfs* and *hfa H. baltica* mutants. Data are expressed as an average of 5 independent replicates and the error bars represent the standard error.

We first deleted the holdfast anchor genes encoding the HfaB and HfaD proteins. Both *H. baltica* Δ*hfaB* and Δ*hfaD* mutants produced holdfast, but they failed to anchor it to the cell body, resulting in holdfast being shed in the medium (Fig. 2A). *H. baltica* Δ*hfaB* was not able to permanently attach to surfaces, and could not form a biofilm (Fig. 2B). In contrast, *H. baltica* Δ*hfaD* mutants were not completely deficient for permanent adhesion, with around 20% of biofilm formed compared to WT (Fig. 2B). These results are in agreement with what was reported for *C. crescentus* Δ*hfaB* and Δ*hfaD* mutants (24), suggesting that the Hfa proteins have a similar function in both organisms.

We then made in-frame deletions of the genes encoding export proteins HfsA and HfsD. These genes are essential for holdfast production in *C. crescentus* (20). Deletion of these genes in *H. baltica* similarly completely abolished holdfast production (Fig. 2A) and surface attachment (Fig. 2B). These results show that deletion of the export genes is sufficient for a complete loss of holdfast production, and that a holdfast is crucial for surface attachment in *H. baltica*.

Finally, we made in-frame deletions of the genes encoding glycosyltransferases HfsG and HfsL, which are essential for holdfast formation in *C. crescentus* (17, 18). Similarly, *H. baltica* Δ*hfsG* and Δ*hfsL* mutants did not produce holdfasts nor form biofilms (Fig. 2A and 2B).

### Effect of modulating *hfsL* and *hfsG* expression of *H. baltica* holdfast properties

We investigated if tunable expression of the *hfsL* and *hfsG* genes, are essential for holdfast production, could change holdfast synthesis and properties. To achieve this goal, we first engineered a replicating plasmid harboring an inducible promoter suitable for *H. baltica*. We adapted the system developed for a tightly controlled heavy metal (copper) promoter inducible system in *Hyphomonas neptunium,* a marine Caulobacterale closely related to *H. baltica* (35). Similarly, we used the promoter for copper resistant protein operon *copAB* (P*cu*) in *H. baltica*) *copA, hbal_0699,* and *copB, hbal_0698*) (Fig. S1A top panel). We first showed that *H. baltica* can tolerate up to 500 µM of CuSO_4_ without significant effect on growth (Fig. S1B-C). We then fused 500 pb upstream of the *copAB* operon (P*cu*) to *lacZ* gene and assembled the construct onto the pMR10 replicating plasmid (Fig. S1A, bottom panel), to assess P*cu* promoter activity using ß-galactosiade as a reporter. We showed that P*cu* is a tightly controlled promoter, with a working inducible range of CuSO_4_ from 10 to 250 µM (Fig. S1D), concentrations that do not impact *H. baltica* growth (Fig S1 B-C).

We expressed *hfsL* or *hfsG* under control of the P*cu* inducible promoter in *H. baltica ΔhfsL* and *ΔhfsG* mutants. In both cases, when the gene expression is highly induced (250 µM CuSO_4_), holdfast size and adhesion are restored to WT levels (Fig. 3A-B). At lower level of induction (10 µM CuSO_4_), both complemented strains produced small holdfasts (Fig. 3A), but failed to form biofilms after 12 h (Fig. 3B). To test if these results were due to altered adhesive properties of these smaller holdfasts, or if their smaller size was not enabling cells to be retained on the surface, we combined the *ΔhfsL* and *ΔhfsG* mutations with an in-frame deletion of the holdfast anchor gene *hfaB*, resulting in mutants that produce holdfasts shed in the medium upon CuSO_4_ induction (Fig. 3C). We grew exponential cultures of the double-mutants on glass coverslips for 4 h to allow them to attach to the surface. After incubation, the slides were rinsed with dH_2_O to remove all cells that are unable to anchor their holdfast to their cell body, resulting in coverslips displaying attached holdfasts and no cells (Fig. 3C). At low level of induction of *hfsL* or *hfsG,* shed holdfasts from *H. baltica ΔhfaB ΔhfsL* and *ΔhfaB ΔhfsG*, though smaller than those from the *H. baltica ΔhfaB,* were still able to efficiently bind to glass slide (Fig. 3C). We quantified the number of holdfasts attached at different levels of induction of *hfsL* and *hfsG* (Fig. 3D). At low induction, the mutants produced 50% the amount of holdfast compared to WT holdfasts (Fig. 3D). To visualize how cells with small holdfasts interact with glass surface, we performed time-lapse microscopy in a microfluidic device, starting with static conditions, and adding flow after 2 minutes, to allow cells to bind to the surface (Fig. 3E). We observed that at low induction of *hfsL* (10 µM CuSO_4_), cells efficiently bind to the surface, despite their small holdfasts. However, once flow was added to the microfluidic device, cells detached, showing that the small holdfasts are not sufficient to withstand high shear forces. At high induction of *hfsL* (250 µM CuSO_4_), cells produced bigger holdfasts and are able to bind to the surface permanently. This result confirms that the smaller holdfasts are still adhesive, but their size is probably not sufficient to allow the cells to efficiently permanently bind to surfaces and form biofilms.

**Figure 3:**
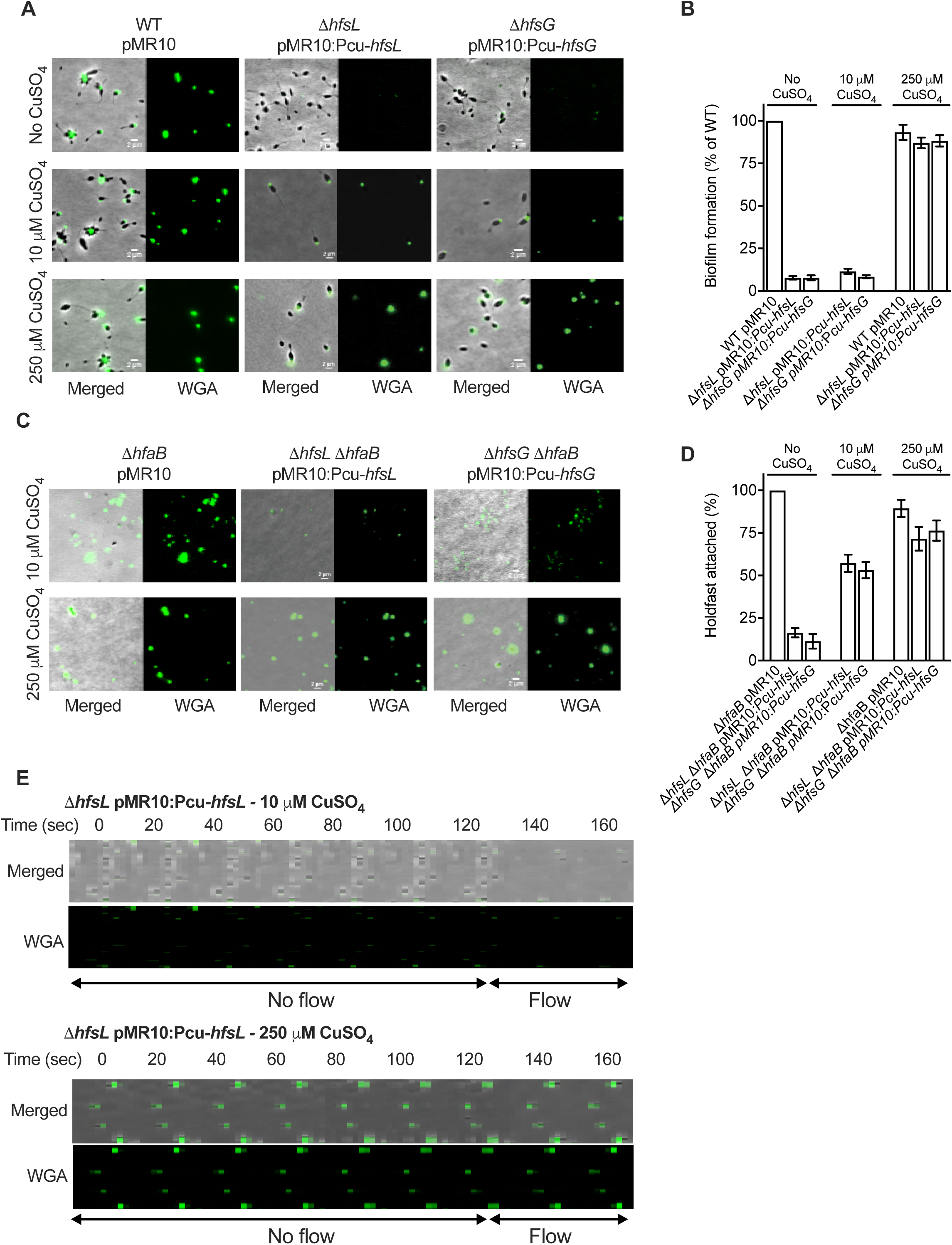
Effect of modulating *hfsL* and *hfsG* expression in *H. baltica* holdfast properties. **A.** Representative images showing merged phase and fluorescence channels of *H. baltica* WT, Δ*hfsL* and Δ*hfsG* mutants complemented with copper inducible promoter constructs and grown in marine broth with 0 µM,10 µM and 250 µM CuSO_4_. Holdfast is labeled with WGA-AF488. **B.** Biofilm quantification after 12 h using crystal violet assay of Δ*hfsL* and Δ*hfsG* mutants and complementations under copper inducible promoter in marine broth supplemented with 0 µM, 10 µM and 250 µM CuSO_4_. Data are expressed as an average of 6 independent replicates and the error bars represent the standard error. **C.** Images of WGA-AF488 labeled *H. baltica* Δ*hfaB, H. baltica* Δ*hfaB* Δ*hfsL* pMR10:P_cu_*hfsL, H. baltica* Δ*hfaB* Δ*hfsG* pMR10:P_cu_*hfsG* shed holdfasts bound to glass slide. Cells were grown in marine broth with 0 µM, 10 µM and 250 µM CuSO_4_ induction for 4 hrs. **D.** Percentage of holdfasts bound to glass slide per field of view at different CuSO_4_ induction measured in (C). Data are expressed as an average of 5 independent replicates and the error bars represent the standard error. **E.** Time-lapse montage of a *H. baltica* Δ*hfsL* pMR10:P_cu_*hfsL* induced with 10 µM (upper panels) and 250 µM (lower panels) CuSO_4_ in a microfluidic device with initially no flow and then low flow introduced to the microfluidic device. Arrows represent time when no flow (first 120 seconds) and flow (later times) was applied to the device.

### *H. baltica* produces large holdfasts by developmental and surface contact stimulation pathways

It was previously shown that WGA interacts with *C. crescentus* and *H. baltica* holdfasts (27, 28). However, side-by-side microscopy imaging using fluorescent WGA suggested that *H. baltica* holdfasts might be larger than *C. crescentus* holdfasts (Fig. 4A). To quantify relative holdfast size, we imaged mixed cultures of *H. baltica* and *C. crescentus* simultaneously labeled with fluorescent WGA lectin. We measured the area of fluorescent WGA staining on single cells for each strain (Fig. 4A) and determined that, on average, the fluorescence area is 5 times larger for *H. baltica* holdfasts compared to those of *C. crescentus* (Fig. 4B). Since WGA binds to GlcNAc residues in the holdfast, either *H. baltica* holdfasts are larger than those of *C. crescentus* or *H. baltica* and *C. crescentus* holdfasts are similar in size, but *H. baltica* holdfasts contain more GlcNAc residues, yielding an increased fluorescence area from bound WGA. To reliably measure the size of holdfasts, we used Atomic Force Microscopy (AFM) and imaged dry holdfasts deposited on a clean mica surface, free of any straining. Results confirmed that *H. baltica* produces larger holdfasts than *C. crescentus*. *H. baltica* holdfasts have a median height of 68 nm, while *C. crescentus* produces holdfasts with median height of 19 nm (Fig. 4C-D), in agreement with previous reports (6, 36).

**Figure 4:**
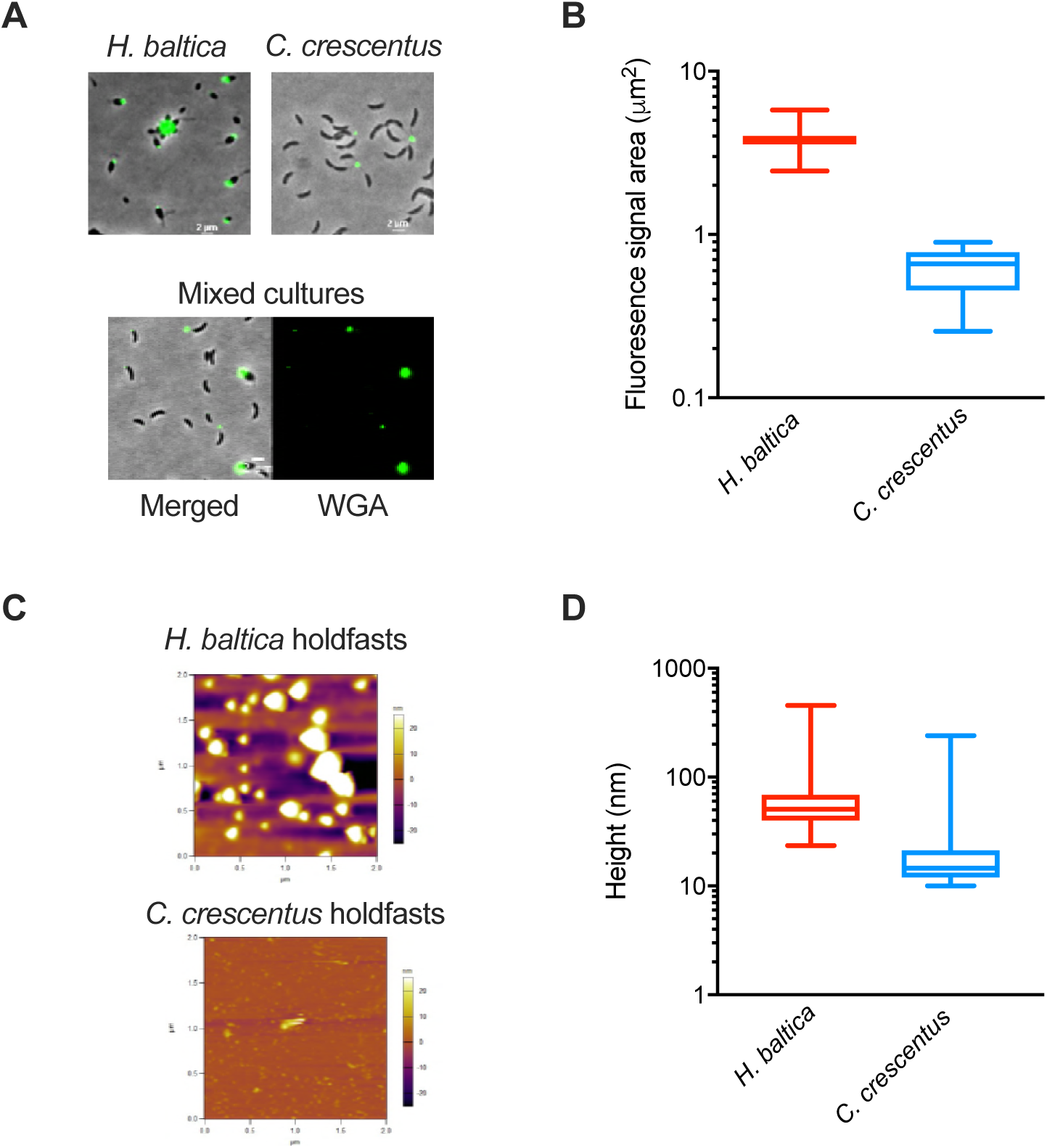
*H. baltica* produces large holdfasts. **A.** Images of *H. baltica, C. crescentus* and mixed culture with holdfasts labeled with WGA-AF488 (green). **B.** Quantification of holdfast size based on fluorescent area covered by WGA-AF488 collected in (A). Data on the box and whiskers plots represent 5 independent replicates of 200 holdfasts from each strain. **C** AFM images of dry shed holdfasts from *H. baltica ΔhfaB* and *C. crescentus ΔhfaB* deposited on a mica surface. The colors on the scale represent the height of the holdfast relative to the surface. **D.** Box and whiskers plots of holdfast height distribution from the AFM images collected in (C). More than 500 holdfasts were measured in 10 independent images.

*C. crescentus* can produce holdfast by two distinct pathways, as part of a complex developmental program in a cell cycle regulated manner or upon contact with a surface, independent of the cell cycle (37-40). Some Alphaproteobacteria, such as *Asticaccaulis biprosthecum* (38) or *Prosthecomicrobium hirschii* (41) are also able to produce holdfasts via developmental and surface-contacted stimulated pathways, while others, like *Agrobacterium tumefaciens*, only produce holdfasts upon contact with a surface (38, 42). To determine how holdfast production is regulated in *H. baltica,* we measured the timing of holdfasts synthesis in the presence or absence of a hard surface. To test whether *H. baltica* holdfast production can be stimulated upon contact with a surface, we performed time-lapse microscopy in a microfluidic device where cells are in close proximity with a glass surface, and we tracked single cells as they reached the surface. We observed holdfast production by including fluorescently labeled WGA in the medium, and we recorded the difference between the time when a cell first reaches the surface and the time when a holdfast is synthesized (Fig. 5A, top panels). We observed that *H. baltica* produces holdfasts within approximately 3 min upon surface contact, (Fig. 5A-B), showing that this species is able to trigger holdfast synthesis upon contact with a surface. To assess cell cycle progression and timing of holdfast synthesis independent of a surface, we tracked single cells and monitored cell differentiation and holdfast synthesis by time-lapse microscopy on soft agarose pads containing fluorescent WGA (Fig 5A, lower panel and 5B). *H. baltica* newborn swarmer cells produced holdfast within 15-25 minutes after budding on an agarose pad (Fig. 5A-B), showing that *H. baltica* can produce holdfasts through progression of the cell cycle, as part of a developmental pathway. To determine the timing of holdfast production relative to the cell cycle length, we measured the time required for a newborn swarmer cell to complete its first and second budding divisions on agarose pads (Fig. 5C). *H. baltica* swarmer cells complete their first budding within 160 - 200 mins (Fig. 5C), similarly to *C. crescentus* production of swarmer cells in PYE complex medium (43), meaning that the holdfast is synthetized within ∼1/10 of the cell cycle, similarly to *C. crescentus* (37).

**Figure 5:**
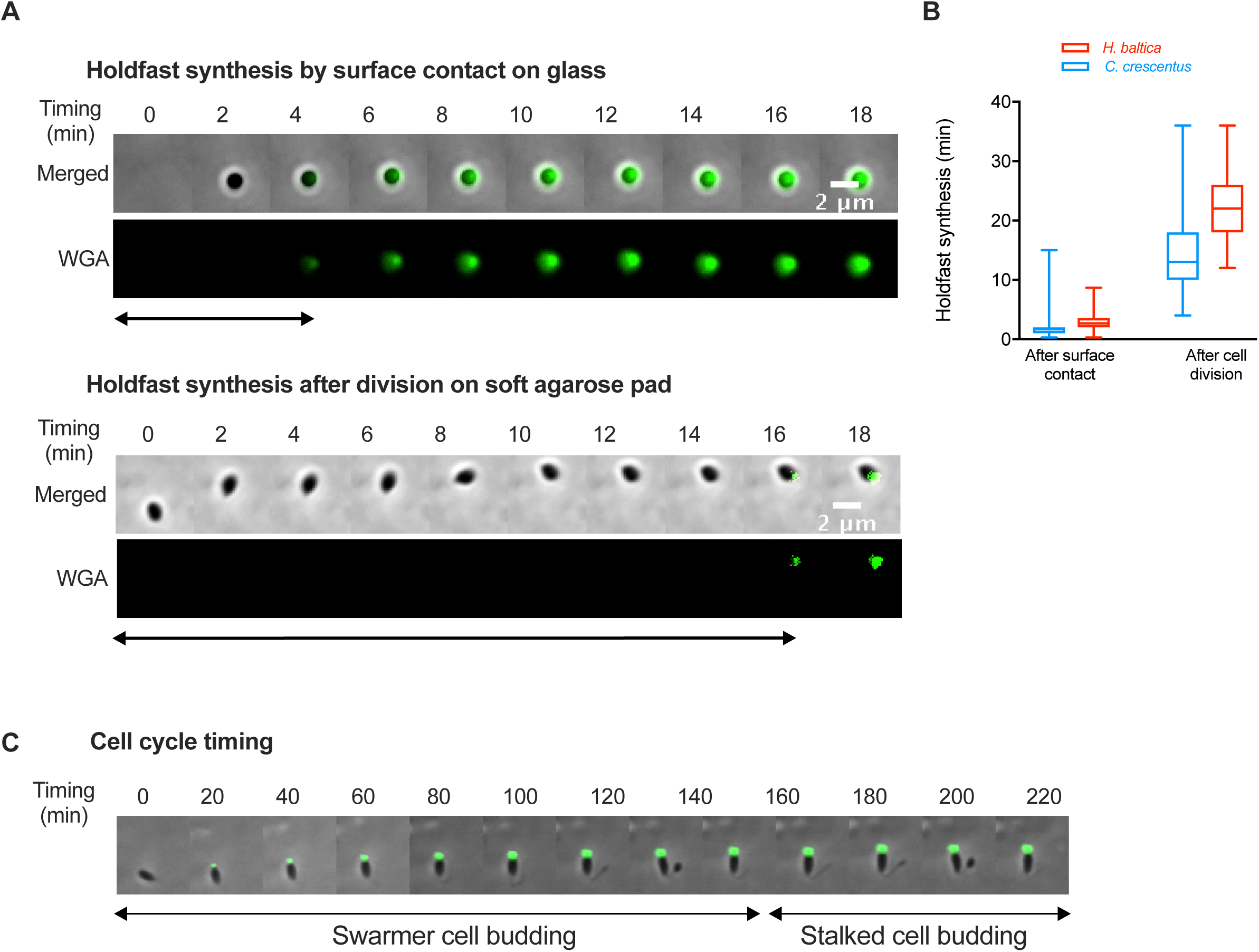
*H. baltica* produces holdfasts via developmental pathway and upon contact with a surface. **A.** Montages of *H. baltica* holdfast synthesis by a newly budded swarmer cell on a glass surface on a microfluidic device (holdfast production after surface contact, top panel), and on soft agarose pads (holdfast production after cell division, bottom panel). Holdfasts are labeled with WGA-AF488 (green). Images were acquired every 20 sec (top panel) and 2 min (bottom panel), and holdfast synthesis timing was processed using MicrobeJ. The arrow indicates the time it takes for holdfast to be detected after surface contact. **B.** Box and whisker plots representing the quantification of *H. baltica* holdfast timing via surface contact stimulation and developmental pathway. Data for *C. crescentus* holdfast synthesis timing were extracted from (2). Total number of cells analyzed is 100 for each set up. **C.** Time-lapse montage of a *H. baltica* swarmer cell differentiating into a budding stalked cell on agarose pad containing WGA-AF488 to label holdfast. Images were collected every 5 min for 3 h. The arrow indicates the time it takes for holdfast to be detected after cell division.

### *H. baltica* holdfast contains GlcNAc and galactose monosaccharides and proteins

Holdfasts in diverse Alphaproteobacteria bind to WGA, showing that they contain GlcNAc residues (3). Previous studies using lectin labelling showed that GlcNAc polymers are the main polysaccharide present in *C. crescentus* holdfast, while other Caulobacterales strains may have additional monosaccharides in their holdfasts (10). Indeed, WGA lectin (specific to GlcNAc) and Dolichos Biflorus agglutinin (specific to N-acetylgalactosamine) both bind *Caulobacter henricii* holdfasts (10), while *Caulobacter subvibriodes* holdfasts was shown to interact with Dolichos Biflorus Agglutinin (specific to N-acetylgalactosamine), Concanavalin A (specific to *α*-mannose) and Ulex Europaeus agglutinin (specific to *α*-fucose), but not WGA (10).

To identify the type of saccharides present in *H. baltica* holdfast, we screened a variety of fluorescent lectins to attempt to label *H. baltica* holdfast (Table 2 and Table S3). Our results indicate that, in addition to binding to WGA, *H. baltica* holdfast also binds to Solanum Tuberosum potato lectin (STL), Lycopersicon Esculentum tomato lectin (LEL), and Datura Stramonium Lectin (DSL1), all lectins specific to GlcNAc residues (Table 2), confirming that *H. baltica* holdfasts contain GlcNAc residues. In addition, lectins that specifically recognize *α*-galactose residues, Griffonia Simplicifolia (GSL1), and Ricinus Communis Agglutinin 1 (RCA120), also interact with *H. baltica* holdfasts (Table 2), while not binding to *C. crescentus* holdfasts (Fig. 6A). Interestingly, Soybean Agglutinin lectin (SBA) did not bind to *H. baltica* holdfasts, showing that these holdfasts only contain galactose and no N-acetylgalactosamine residues (GalNAc) (Table 2). These results show that *H. baltica* holdfasts have a different sugar composition than Caulobacter and contain both GlcNAc and galactose residues. To confirm that observed galactose-specific binding was holdfast dependent, we labeled *H. baltica* Δ*hfsA,* Δ*hfsG* (holdfast minus strains) and *ΔhfaB* (holdfast shedding strain) mutants with both WGA and GSL1 lectins. None of the lectins labelled the holdfast deficient Δ*hfsA* and *ΔhfsG* mutants, but they labelled shed holdfast produced by the Δ*hfaB* mutant (Fig. 6A), confirming that *H. baltica* holdfasts contain galactose residues.

**Table 2:** Lectin binding assays

| Lectin | Specificity | *H. baltica*<br>*holdfast* | *C. crescentus*<br>*holdfast* |
| --- | --- | --- | --- |
| Wheat Germ Agglutinin | GlcNAc | √ | √ |
| Lycopersicon Esculentum Tomato Lectin | GlcNAc 1-4 | √ | √* |
| Datura Stramonium Lectin | GlcNAc 1-4 | √* | - |
| Solanum Tuberosum Potato Lectin | GlcNAc , prefers trimers and tetramers | √ | √* |
| Ricinus Communis Agglutinin | Galactose | √ | - |
| Griffonia Simplicifolia Lectin 1 | α-GalNAc, α-galactose | √ | - |
| Soybean Agglutinin | α-GalNAc | - | - |

**Figure 6:**
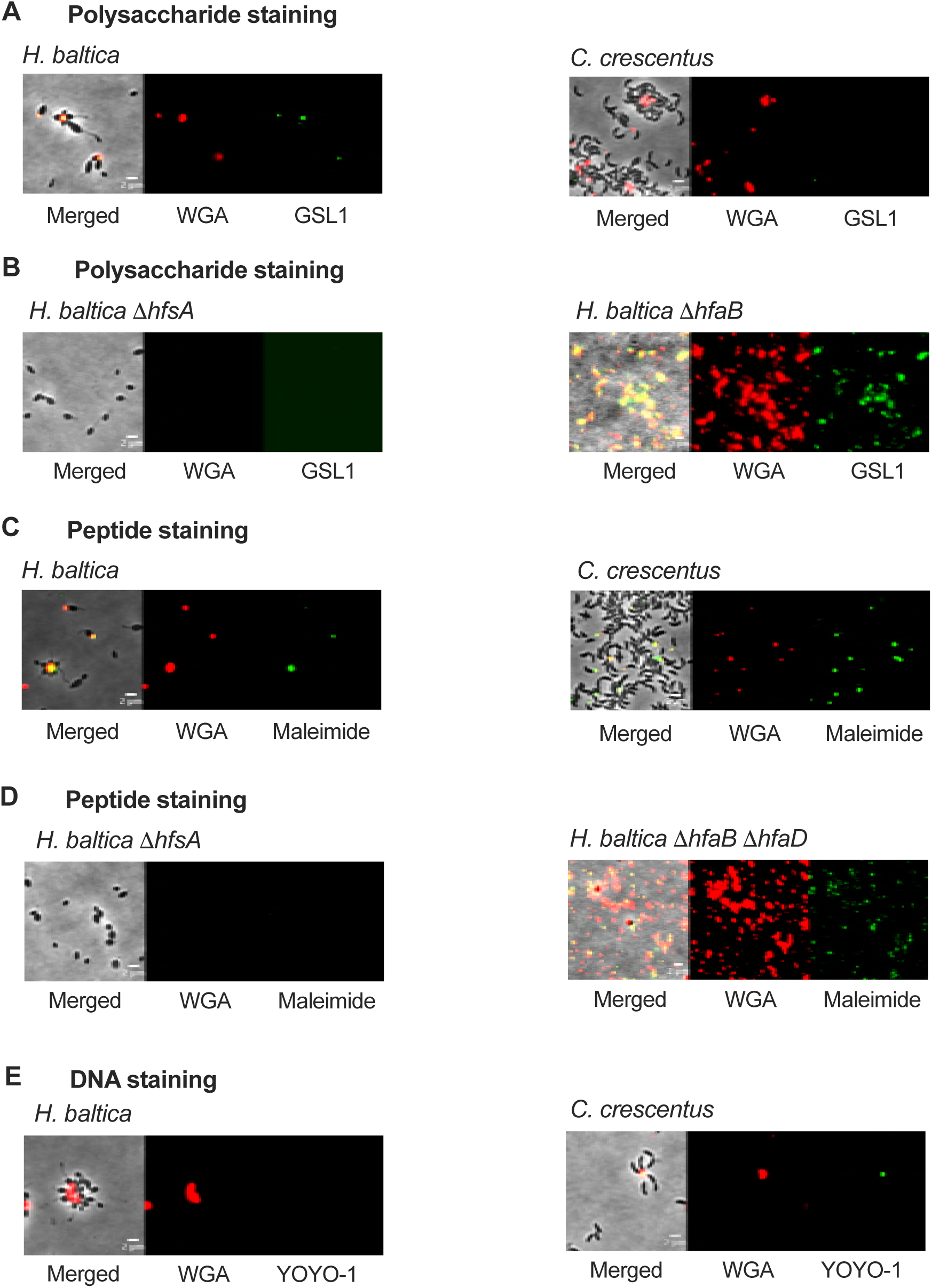
*H. baltica* holdfast contains GlcNAc and galactose monosaccharides, and proteins. **(A-C)** Representative images showing merged phase and fluorescence channels on the left and fluorescence channels alone on the middle and right. **A.** *H. baltica* and *C. crescentus* holdfasts were co-labeled with WGA-AF594 (GlcNAc) and GSL1-AF488 (galactose) lectins, to stain polysaccharides. **B** *H. baltica* and *C. crescentus* holdfasts were co-labeled with WGA-AF594 (GlcNAc) lectin and AF488mal, to stain peptides. **C.** *H. baltica* and *C. crescentus* holdfasts were co-labeled with WGA-AF594 (GlcNAc) lectin and YOYO-1-AF488, to stain DNA.

*C. crescentus* holdfasts have been recently shown to contain peptides and DNA residues (15). To test whether *H. baltica* holdfast contains proteins, we attempted to label putative cysteines in the holdfast using a fluorescent maleimide dye (AF488mal). As for *C. crescentus* holdfasts, *H. baltica* holdfasts could be stained with AF488mal, showing that these holdfasts possess molecules with free accessible thiols, suggesting the presence of peptides containing cysteines (Fig. 6B). The staining was holdfast-specific, as AF488mal did not label the holdfast-deficient Δ*hfsA* and Δ*hfsG* mutants (Fig. 6B). It has been shown that in *C. crescentus*, holdfast labeling by AF488mal was specific to holdfasts attached to cells, as shed holdfasts from a holdfast anchor mutant were not labeled, suggesting that the cysteine-containing HfaD in cell-anchored holdfasts is responsible for the labeling of those holdfasts with AF488mal (15). In *H. baltica*, both the anchor proteins HfaB and HfaD contain cysteines. In order to test whether AF488mal interacts with HfaB or HfaD, we stained shed holdfasts produced by a *H. baltica* Δ*hfaB ΔhfaD* double mutant and could detect staining (Fig. 6B). This is in stark contrast with *C. crescentus* holdfasts that react with AF488mal only when attached to WT cells (15, 44). This result show that the holdfast composition in these two microorganisms is different.

To probe for the presence of DNA in *H. baltica* holdfasts, we labeled holdfasts with the fluorescent DNA dye YOYO-1 that binds to double-stranded DNA molecules. As previously reported, *C. crescentus* holdfasts was labeled with YOYO-1 (15). However, YOYO-1 failed to label *H. baltica* holdfasts (Fig. 6C), suggesting that *H. baltica* holdfasts do not contain DNA. It has been previously shown that, in *C. crescentus,* extracellular DNA (eDNA) released during *C. crescentus* cell lysis binds specifically to *C. crescentus* holdfasts, preventing adhesion to surfaces and biofilm formation (45), and it has been hypothesized that it could be due to a specific interaction between the DNA present in the holdfast and eDNA (15). We showed above that *H. baltica* holdfasts were devoid of DNA, so we tested whether eDNA could inhibit *H. baltica* binding. We performed short term adhesion assays in the presence of *H. baltica* and *C.crescentus* eDNA (Fig. S2A). When *C. crescentus* eDNA is present, the number of *C. crescentus* attached to the glass slide after 60 minutes is dramatically decreased, compared to when *H. baltica* eDNA is added and to the no DNA addition control (Fig S2A), confirming previous studies that showed that, in *C. crescentus,* eDNA inhibition was specific for *C. crescentus* eDNA (45). However, *H. baltica* adhesion is not impaired by the presence of eDNA, from itself or from *C. crescentus* (Fig. S2A). We also performed long term biofilm assays in the presence of eDNA and showed that *H. baltica* biofilm formation is not impaired by the presence of eDNA in the medium after 24 h of incubation (Fig. S2B).

Taken together, we show that *H. baltica* holdfasts are different from *C. crescentus* ones: they are larger, contain GlcNAc, galactose, and peptide residues, but are void of DNA.

### *H. baltica* holdfast tolerates high ionic strength

It has been shown that *C. crescentus* holdfasts are very sensitive to ionic strength, as purified holdfast binding efficiency to glass decreased by 50% with addition of 10 mM NaCl (6). *C. crescentus* is a freshwater bacterium and has probably evolved without selective pressure to bind under high ionic strength. This compelled us to investigate how the holdfasts from *H. baltica* are affected by ionic strength. We first used NaCl to study the effects of ionic strength on holdfast binding, since it is the most abundant ionic elements in marine water and it has been used in many studies to assess the effect of ionic strength on bacterial adhesins (6, 46-48). We quantified purified holdfast binding to glass at different NaCl concentrations, using fluorescent WGA, and plotted the relative number of holdfasts per field of view bound to glass at different concentrations of NaCl (Fig. 7A-B). Our results confirmed that *C. crescentus* holdfast is very sensitive to NaCl, as only 50% of holdfasts can bind to glass when 10 mM NaCl is added (Fig. 7B). However, *H. baltica* holdfast tolerated up to 500 mM NaCl without any effect on surface binding (Fig. 7B). There was a 50% decrease in *H. baltica* holdfast binding at 600 mM (Fig. 7B), showing that *H. baltica* holdfasts are more than 50 times more resistant to NaCl than those of *C. crescentus*. *H. baltica* was originally isolated from the Baltic Sea, which has 250 mM NaCl (Fig 7B, gray arrow) (26), and at that NaCl concentration, the binding efficiency of *H. baltica* holdfasts is maximal. Interestingly, *H. baltica* holdfasts still bound efficiently at low ionic strength. We observed similar results using different concentrations of MgSO_4_ (Fig. 7C): *H. baltica* holdfasts were 50 times more resistant to MgSO_4_ than those of *C. crescentus*, showing that the binding inhibition is not specific to NaCl but is rather dependent on ionic strength.

**Figure 7:**
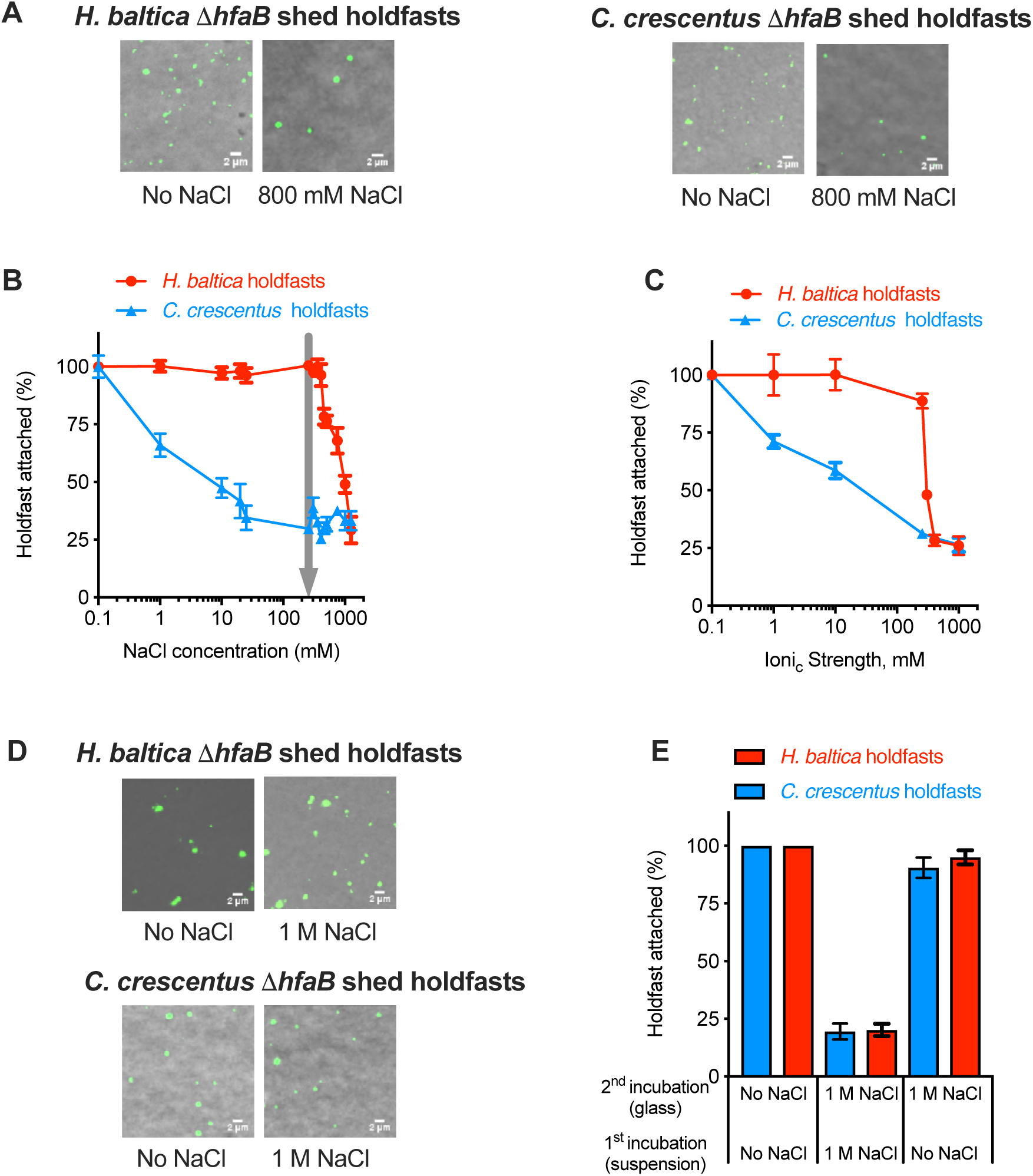
*H. baltica* holdfast tolerates higher ionic strength. **A.** Images of WGA-AF488 labeled *H. baltica ΔhfaB* and *C. crescentus ΔhfaB* shed holdfasts bound to glass slide, incubated in different concentration of NaCl for 4 hours. **B.** Percentage of holdfasts bound per field of view at different concentrations of NaCl. Gray arrow indicates ionic strength of marine broth and Baltic sea (250 mM), natural habitat for *H. baltica*. Data are expressed as an average of 6 independent replicates and the error bars represent the standard error. **C.** Percentage of holdfasts bound per field of view at different concentrations of MgSO_4_. Data are expressed as an average of 4 independent replicates and the error bars represent the standard error **D.** Images of WGA-AF488 labeled holdfasts already bound to a glass surface and incubated in 0 mM NaCl (left) and 1 M NaCl (right) for 12hrs. **E.** Percentage of holdfasts bound per field of view at 0 M or 1 M NaCl. The first incubation was done by adding 0 M or 1 M of NaCl to a holdfast suspension spotted on a glass slide. After a 12 h incubation, the second incubation was done after washing off unbound holdfasts, by adding 0 M or 1 M NaCl directly to the holdfasts attached to the glass slide, and by incubating for another 12 h. Data are expressed as an average of 5 independent replicates and the error bars represent the standard error.

Our results show that, in *H. baltica*, initial holdfast binding to glass is not changed for NaCl concentrations up to 500 mM, then drastically decreased to reach around 25% of holdfasts attached at 1 M NaCl (Fig. 7B). To test whether high ionic strength could remove holdfasts previously attached to the glass surface, we first incubated purified holdfasts for 4 h without any salt added, and then added 1M of NaCl for 12 hours to the bound holdfasts (Fig. 7D). Bound holdfasts from *H. baltica* and *C. crescentus* were not dislodged from the glass surface (Fig. 7D and E), indicating that while high ionic strength inhibits holdfast from binding to the surface, it cannot dislodge bound holdfast from a glass surface (Fig. 7E).

## DISSCUSION

Different bacterial species harbor an adhesive holdfast and use it to attach to surfaces (2, 3, 9, 49, 50). They represent an extremely diverse group in terms of their physiology and the natural environments they inhabit (soil, freshwater, and marine environments). They have evolved the ability to adhere to surfaces with vastly different composition in varying environmental conditions (salinity, pH, temperature, etc.). Holdfast chemical properties have been mainly studied in the model organism *C. crescentus,* a freshwater Caulobacterale (6, 10, 13, 15, 18, 28, 32, 36, 51), and little is known about holdfast properties and composition in marine Caulobacterales. In this study, we used *H. baltica* as a model species living in a marine environment and found that this bacterium has a holdfast tailored for adhesion in high salinity conditions. We show that holdfasts in *H. baltica* are different than those of *C. crescentus*: they are larger, have a different chemical composition, and have a high tolerance to ionic strength.

The bioinformatics analysis of holdfast genes indicated that the *hfs* and *hfa* loci are highly conserved among Caulobacterales, with few reshufflings of these genes (Fig. 1C). The arrangements of the holdfast genes in the *hfs* and *hfa* loci appears to be ancestral while the relocation of some of the genes is a recent event that could affect their level of expression (52). Through deletion and complementation of important *hfs* and *hfa* genes, we confirmed that holdfast biogenesis and anchoring to the cell body in *H. baltica* use similar genes to those identified in *C. crescentus* (2);(18) (Fig 2).

We showed that the two glycosyltransferases *hfsL* and *hfsG,* are essential for holdfast production and regulate the amount of sugar monosaccharides added to holdfast polysaccharides, as cells expressing low levels of these proteins produce smaller holdfasts (Fig. 2, 3). Small holdfasts with less polysaccharides binds to glass but not strongly enough to support cells (Fig. 3). This phenomenon could be due to the smaller surface contact area between the small holdfasts being insufficient to resist drag and shear forces during the washing steps of our assays or to a change in holdfast structure or composition due to the lower expression of the glycosyltransferases HfsL and HfsG. More studies on the role of HfsL and HfsG will help us to determine if these enzymes play an important role in specific physicochemical properties of *H. baltica* holdfasts.

In *C. crescentus*, the growing holdfast polysaccharide repeat units are modified by the acetyltransferase HfsK (32) and the polysaccharide deacetylase HfsH (21) (Fig. 1A). These two enzymes are not essential for holdfast production in *C. crescentus*, but modify adhesiveness and cohesiveness of the holdfast. Holdfasts produced by Δ*hfsH* or Δ*hfsK* mutants produced thread-like holdfasts with weaker adhesion strength (28, 32). In addition, fully acetylated purified holdfasts from the *C. crescentus* Δ*hfsH* mutant holdfasts were not affected by ionic strength (6), suggesting that holdfast modification can modulate salt tolerance. Our future work will determine how holdfast modification impacts *H. baltica* holdfasts tolerance to high ionic, and the possible role of HfsH and HfsK.

The exact composition and structure of holdfast is still unknown in the model organism *C. crescentus*. Lectin binding assays and lysozyme treatment support GlcNAc as one of the important components in holdfasts (10, 36). Treating holdfast with proteinase K and DNase I affects *C. crescentus* holdfast structure and force of adhesion, suggesting that it contains peptide and DNA residues (15). In this work, we identified different components present in *H. baltica* holdfasts: these holdfasts contain galactose monosaccharides in addition to GlcNAc (Fig. 6A). In the different *hfs* mutants generated in this study, galactose monosaccharides were not detected on the cell pole (Fig. 6A), suggesting that GlcNAc and galactose are produced together or secreted by the same proteins. Shed holdfasts from *H. baltica ΔhfaB* contain both GlcNAc and galactose (Fig. 6A), implying that they are both anchored to the cell envelope with the same anchor proteins. *H. baltica* holdfasts are void of DNA, a stark contrast to *C. crescentus* (Fig. 6C). In addition, *H. baltica* holdfasts could be successfully stained with a fluorescent maleimide dye, which suggest the presence of a protein or peptide with a cysteine residue (53). The maleimide dye stains only cells with a holdfast, and interacts with holdfasts without the presence of cells, indicating that the reactive molecules are intrinsic part of *H. baltica* holdfast (Fig. 6B), another notable difference with *C. crescentus* holdfasts where maleimide dye only interacts with holdfasts attached to cells (15). Our results suggest that the two holdfasts from *H. baltica* and *C. crescentus* have different composition.

Bacterial adhesins have been shown to use electrostatic and hydrophobic interactions to attach to surfaces (6). Electrostatic interactions are impaired in high ionic environment like seawater with 600 mM of NaCl (7). *C. crescentus* holdfast uses both ionic and hydrophobic interactions and its binding is impaired in presence of NaCl in the media (6). We have shown that *H. baltica* holdfasts tolerate high ionic strength compared to *C. crescentus* (Fig. 7A-C). Marine Caulobacterales face a higher ionic strength environment than the freshwater bacteria, therefore, it is vital that marine Caulobacterales produce holdfasts that are more tolerant to ionic strength and strongly adhere in saline environments. Holdfasts do not efficiently bind at 1 M NaCl, but holdfasts already attached to a surface cannot be removed when adding 1 M NaCl (Fig. 7D), suggesting that the binding inhibition at 1 M NaCl takes place during the initial stage of surface interaction, because it has no effect on surface bound holdfasts (Fig. 7D). These results imply that holdfast interacts with surfaces initially using electrostatic interactions, before a permanent molecular bond is formed (6, 54). The differences in ionic tolerance between fresh and marine Caulobacterales indicates that there are significant differences in physicochemical properties between the two types of holdfasts. Holdfast structure and binding properties could depend on the type and the amount of sugars polymerized in the holdfast polysaccharide that are specialized to interact with different surfaces (55).

In conclusion, we have shown that *H. baltica* produces holdfasts with different binding and physicochemical properties compared to *C. crescentus* holdfasts. This could suggest that there are additional holdfast related genes or regulators that have not been identified. A careful genetic screen in *H. baltica* will provide more insights about holdfast production and the underlying mechanisms yielding to an enhanced adhesion at high ionic strength. The molecular mechanism by which *H. baltica* and other marine Caulobacterales overcome the effect of ionic strength on holdfast binding will be our next focus.

## MATERIALS AND METHODS

### Identification of orthologous holdfast genes and phylogenetic analysis

*C. crescentus* holdfast genes were used to find bi-directional best hits (BBH) on Caulobacterales genomes. The putative genes were selected for E^-^ value > 10^-4^ and sequence identify > 30%. The phylogenic tree was built using 16S rRNA sequences of the selected Caulobacterales. Sequences were aligned using MUSCLE software (46). The aligned sequences were used to construct the maximum likelihood phylogeny using the MEGA6 software (58). The LG+G+I models and analysis of 1000 bootstraps were used to generate the nodes values for each clade.

### Bacterial strains and growth conditions

The bacterial strains used in this study are listed in Table S1. *H. baltica* strains were grown in marine medium (Difco Marine Broth/Agar reference 2216), except when studying the effect of ionic strength on holdfast binding where they were grown in Peptone Yeast Extract (PYE) medium (8) supplemented with 0 or 1.5% NaCl or MgSO_4_. *C. crescentus* was grown in PYE medium. Both *H. baltica* and *C. crescentus* strains were grown at 30 °C. When appropriate, antibiotics were added to the following concentrations: kanamycin (Kan) 5 µg/ml in liquid and 20 µg/ml on agarose plates. *H. baltica* strains with copper inducible promoter were grown in marine broth supplemented with 0-250 µM of CuSO_4_. *E. coli* strains were grown in Luria-Bertani medium (LB) at 37 °C with no antibiotics or with 30 µg/ml of Kan in liquid or 25 µg/ml on agarose plate when needed.

### Strains construction

All the plasmids and primers used in this study are listed on Table S1 and S2 respectively. In-frame deletion mutants were obtained by double homologous recombination as previously described (59), using suicide plasmids transformed into the *H. baltica* host strains by mating or electroporation (60). Briefly, genomic DNA was used as the template to PCR-amplify 500 bp fragments from upstream and downstream regions of the gene to be deleted. pNPTS139 plasmid was cut using EcoRV-HF endonuclease from New England Biolabs. The primers used to amplify 500 bp upstream and downstream of the gene were designed to have overlapping 25 bp for isothermal assembly (61) using the New England Biolabs (NEB) NEBuilder tools for Gibson assembly into plasmid pNPTS139. Then pNPTS139-based constructs were transformed into *α*-select *E. coli* strain and introduced in the host *H. baltica* by mating or electroporation (62). The two-step selection for homologous recombination was carried out using sucrose resistance and kanamycin sensitivity (63).

For gene complementation, the pMR10 plasmid was cut with EcoRV-HF and 500 bp of the promoter and the gene were ligated into plasmid pMR10 using NEBuilder tools. The pMR10-based constructs were transformed into *α*-select *E. coli* strain and introduced in the host *H. baltica* by mating or electroporation, followed by Kan selection. The plasmid constructs and mutants were confirmed by sequencing.

### Holdfast labeling using fluorescently labeled lectins

Alexa Fluor (AF) conjugated lectins (Vector Labs, Table 2 and Table S3) were added to 100 µl of exponential culture to a final concentration of 0.5 µg/ml and incubated at room temperature for 5 min. 3 µl of the labeled culture was spotted on glass cover slide and covered with 1.5 % (w/v) SeaKem LE agarose (Lonza) pad in water and visualized by epifluorescence microscopy. Holdfasts were imaged by epifluorescence microscopy using an inverted Nikon Ti-E microscope with a Plan Apo 60X objective, a GFP/DsRed filter cube, an Andor iXon3 DU885 EM CCD camera and Nikon NIS Elements imaging software with 200 ms exposure time. Images were processed in ImageJ (64).

### Short-term and biofilm binding assays

This assay was performed as previously described (28) with the following modification. For short-term binding, exponential cultures (OD_600_ = 0.6 - 0.8) were diluted to OD_600_ = 0.4 in fresh marine broth, added into 24-well plate (1 ml per well), and incubated shaking (100 rpm) at room temperature for 4 h. For biofilm assays, overnight cultures were diluted to OD_600_ = 0.10, added to 24-well plate (1 ml per well), and incubated at room temperature for 12 hours with shaking (100 rpm). In both set-ups, OD_600_ were measured before the wells were rinsed with distilled H_2_O to remove non-attached bacteria, stained using 0.1% crystal violet (CV) and rinsed again with dH_2_O to remove excess CV. The CV was dissolved into 10% (v/v) acetic acid and quantified by measuring the absorbance at 600 nm (A_600_). The biofilm formation was normalized to A_600_ / OD_600_ and expressed as a percentage of WT.

### *hfsL* and *hfsG* expression using copper inducible promoter

Strains bearing copper inducible plasmids were inoculated from freshly grown colonies into 5 ml marine broth containing 5 μg/ml Kan and incubated shaking (200 rpm) at 30°C overnight. Overnight cultures were diluted in the same culture medium to OD_600_ = 0.10 and incubated until OD_600_ = 0.4 was reached. When needed, copper sulfate dissolved in marine broth was added to a final concentration of 0-250 µM. The induced cultures and controls were added to 24-well plate (1 ml per well) and incubated shaking (100rpm) at room temperature for 4-8 h. Then, OD_600_ were measured before the wells were rinsed with distilled H_2_O to remove non-attached bacteria, stained using 0.1% crystal violet (CV) and rinsed again with dH_2_O to remove excess CV. The CV was dissolved into 10% (v/v) acetic acid and quantified by measuring the absorbance at 600 nm (A_600_). The biofilm formation was normalized to A_600_ / OD_600_ and expressed as a percentage of WT.

### Visualization of holdfasts attached on a glass surface

Visualization of holdfast binding to glass surfaces were performed as described previously in (28) with the following modification. *H. baltica and C. crescentus* strains grown to exponential phase (OD_600_ = 0.2 – 0.6) were incubated on washed glass coverslips at room temperature in a saturated humidity chamber for 4 - 8 h. After incubation, the slides were rinsed with dH_2_O to remove unbound cells, and holdfasts were labelled using 50 µl of fluorescent Alexa Fluor (AF488 or AF594) conjugated lectins (Molecular Probes or Vector Labs, Table 2) at a final concentration of 0.5 µg/ml. Then, slides were rinsed with dH_2_O and topped with a glass coverslip. Holdfasts were imaged by epifluorescence microscopy using an inverted Nikon Ti-E microscope with a Plan Apo 60X objective, a GFP/DsRed filter cube, an Andor iXon3 DU885 EM CCD camera and Nikon NIS Elements imaging software with 200 ms exposure time.

Images were processed in ImageJ (64).

### Atomic Force Microscopy (AFM)

AFM imaging was performed using the tapping mode on a Cypher AFM (Asylum Research) at 20°C, as described previously (6, 19) with the following modifications. Exponential phase grown *H. baltica ΔhfaB* and *C. crescentus ΔhfaB* were diluted and spotted on a freshly cleaved mica. Samples were grown overnight at room temperature in a humid chamber. The samples were then rinsed with sterile dH_2_O to remove unbound cells and debris, and air-dried. AFM topographic images of dried holdfasts attached to the mica surface were obtained using a silicon Olympus AC160TS cantilever (Resonance frequency = 300 kHz, Spring constant = 26 N/m). 40 images of 4 independent replicates were obtained. Holdfast height was determined using the built-in image analysis function of the Igor Pro/Asylum Research AFM software.

### Holdfast synthesis timing by time-lapse microscopy on agarose pads

*H. baltica* holdfast synthesis timing were observed in live cells on agarose pads by time-lapse microscopy as described previously (39) with some modifications. A 1 µl aliquot of exponential-phase cells (OD_600_ = 0.4 – 0.8) was placed on top of a pad containing 0.8% agarose in marine broth with 0.5 µg/ml AF-WGA 488. A coverslip was placed on top of the agarose pad and sealed with VALAP (Vaseline, lanolin and paraffin wax). Time-lapse microscopy images were taken every 2 min for 4 h using an inverted Nikon Ti-E microscope and a Plan Apo 60X objective, a GFP/DsRed filter cube, and an Andor iXon3 DU885 EM CCD camera. Time-lapse movies were visualized in ImageJ (64) to manually assess the timing of a swarmer cell producing holdfast (lectin detection) after budding. The time difference between holdfast synthesis and budding was determined using MicrobeJ (65).

### Holdfast synthesis timing by time-lapse microscopy on microfluidic device

This experiment was performed as previously described (40) with the following modifications. Cell cultures were grown to mid-exponential phase (OD_600_ = 0.4 - 0.6) and 200 µl of culture was diluted into 800 µl fresh marine broth in the presence of 0.5 µg/ml AF-WGA 488 for holdfast labeling. One ml of the cell culture was flushed into a microfluidic device containing a 10 µm high linear chamber fabricated in PDMS (Polydimethylsiloxane) as described previously (39). After injection of the cells into the microfluidic chamber, the flow rate was adjusted so that attachment could be observed under static conditions or low flow rate.

Time-lapse microscopy was performed using an inverted Nikon Ti-E microscope and a Plan Apo 60X objective, a GFP/DsRed filter cube, an Andor iXon3 DU885 EM CCD camera and Nikon NIS Elements imaging software. Time-lapse videos were collected for strains over a period of 3 h at 20-second intervals. Cell attachment was detected at the glass-liquid interface within the microfluidic chamber using phase contrast microscopy while holdfast synthesis was detected using fluorescence microscopy. Cells that hit the surface and attached permanently via their holdfast during this 3 h period were analyzed for the timing of holdfast synthesis. The time difference between holdfast synthesis and cell-surface contact was determined using MicrobeJ (65) and define as holdfast delay. Cells that were present on the surface at the start of the time-lapse experiment were not analyzed.

### Holdfast labeling using fluorescently labeled Maleimide and YOYO-1

Alexa Fluor (AF-mal488) conjugated Maleimide C_5_ (ThermoFisher Scientific) were added to 100 µl of exponential culture to a final concentration of 0.5 µg/ml and incubated at room temperature for 5 mins. Similarly, YOYO-1 (fluorescent DNA stain, Molecular Probes) was added to 100ul of exponential culture to a final concentration of 0.5 µg/ml and incubated at room temperature for 5 mins. 3 µl of the labeled culture was spotted on glass cover slide and covered with 1.5 % (w/v) agarose pad in water and visualized by epifluorescence microscopy. Holdfasts were imaged by epifluorescence microscopy using an inverted Nikon Ti-E microscope with a Plan Apo 60X objective, a GFP/DsRed filter cube, an Andor iXon3 DU885 EM CCD camera and Nikon NIS Elements imaging software with 200 ms exposure time. Images were processed in ImageJ (64).

### Effect of ionic strength on holdfast binding

Purified holdfasts attached to a surface in different ionic strength were visualized as described previously (Berne *et al*., 2013), with few modifications. Briefly, *H. baltica ΔhfaB* and *C. crescentus* Δ*hfaB* cells were grown to late exponential phase (OD_600_ = 0.6 – 0.8) in PYE + 1.5% NaCl, and plain PYE respectively. The cells were pelleted by centrifugation for 30 min at 4,000 x *g* and resuspended in PYE and incubated for 2 h at 30 °C. Then, the cells were again pelleted by centrifugation and 100 µl of supernatant, containing free holdfasts shed by the cells, were mixed with 100 µl of NaCl in PYE to make a final concentration of 0 - 1000 mM of NaCl. 50 µl of the mixture was incubated on washed glass coverslips at room temperature in a saturated humidity chamber for 4 - 12 h. After incubation, the slides were rinsed with dH_2_O to remove unbound material, and labelled holdfast were visualized with Alexa Fluor lectins (Vector labs). Holdfasts were imaged by epifluorescence microscopy using an inverted Nikon Ti-E microscope with a Plan Apo 60X objective, a GFP/DsRed filter cube, an Andor iXon3 DU885 EM CCD camera and Nikon NIS Elements imaging software with 200 ms exposure time. Images were processed in ImageJ (64). The number of holdfasts bound per field of view was quantified using MicrobeJ (65).

### β-galactosidase assays to assess copper inducible *copA* promoter activity

This assay was performed as previously described with few modification (44). Strains bearing plasmids with *lacZ* gene controlled by copper inducible promoter *copAB* were inoculated from freshly grown colonies into 5 ml marine broth containing 5 μg/ml Kan and incubated at 30°C overnight. Overnight cultures were diluted in the same culture medium to OD_600_ = 0.10 and incubated until an OD_600_ = 0.4 was reached, where copper sulfate dissolved in marine broth was added to a final concentration of 0 - 250 µM. The induced cultures and controls were incubated for 2 - 4 h at 30 °C. β-galactosidase activity was measured colorimetrically as described previously (66). Briefly, 200 µl of culture was mixed with 600 µl Z buffer (60 mM Na_2_HPO_4_, 40 mM NaH_2_PO_4_, 10 mM KCl, 1 mM MgSO_4_, 50 mM ß- mercaptoethanol). Cells were then permeabilized using 50 µl chloroform and 25 µl 0.1% SDS. 200 µl of substrate *o*-nitrophenyl-β-D-galactoside (4 mg/ml) was added to the permeabilized cells. Upon development of a yellow color, the reaction was stopped by raising the pH to 11 with addition of 400µl of 1M Na_2_CO_3_. Absorbance at 420 nm (A_420_) was determined and the Miller Units of β-galactosidase activity were calculated as (A_420_)(1000)/(OD_600_)(*t*)(*v*) where *t* is the time in minutes and *v* is the volume of culture used in the assay in mL.

### Growth measurements

Impact of CuSO_4_ on *H. baltica* growth was measured using 24-well plates. 1 ml of cultures (Starting OD_600_ = 0.05) were incubated for 12 h at 30°C, using marine Broth and various CuSO_4_ concentrations. OD_600_ were recorded after overnight incubation, to determine the growth yield for the different CuSO_4_ concentrations. Growth curves using 0 or 500 µM CuSO_4_ were recorded every 30 minutes for 20 h. All OD_600_ were recorded using a Biotek Synergy HT.

## ACKNOWLEDGEMENTS

We thank Bogdan Dragnea, Department of Chemistry, Indiana University, for use of his AFM and facilities for analysis of the shed holdfasts. We thank the members of the Brun laboratory for the comments on the manuscript. This work was supported by National Institute of Health Grant R01GM102841 and R35GM122556 to Y.V.B. and a fellowship from the Department of Biology, Indiana University to NKC. Y.V.B holds a Canada 150 Research Chair in Bacterial Cell Biology.

---

**Table S1:**
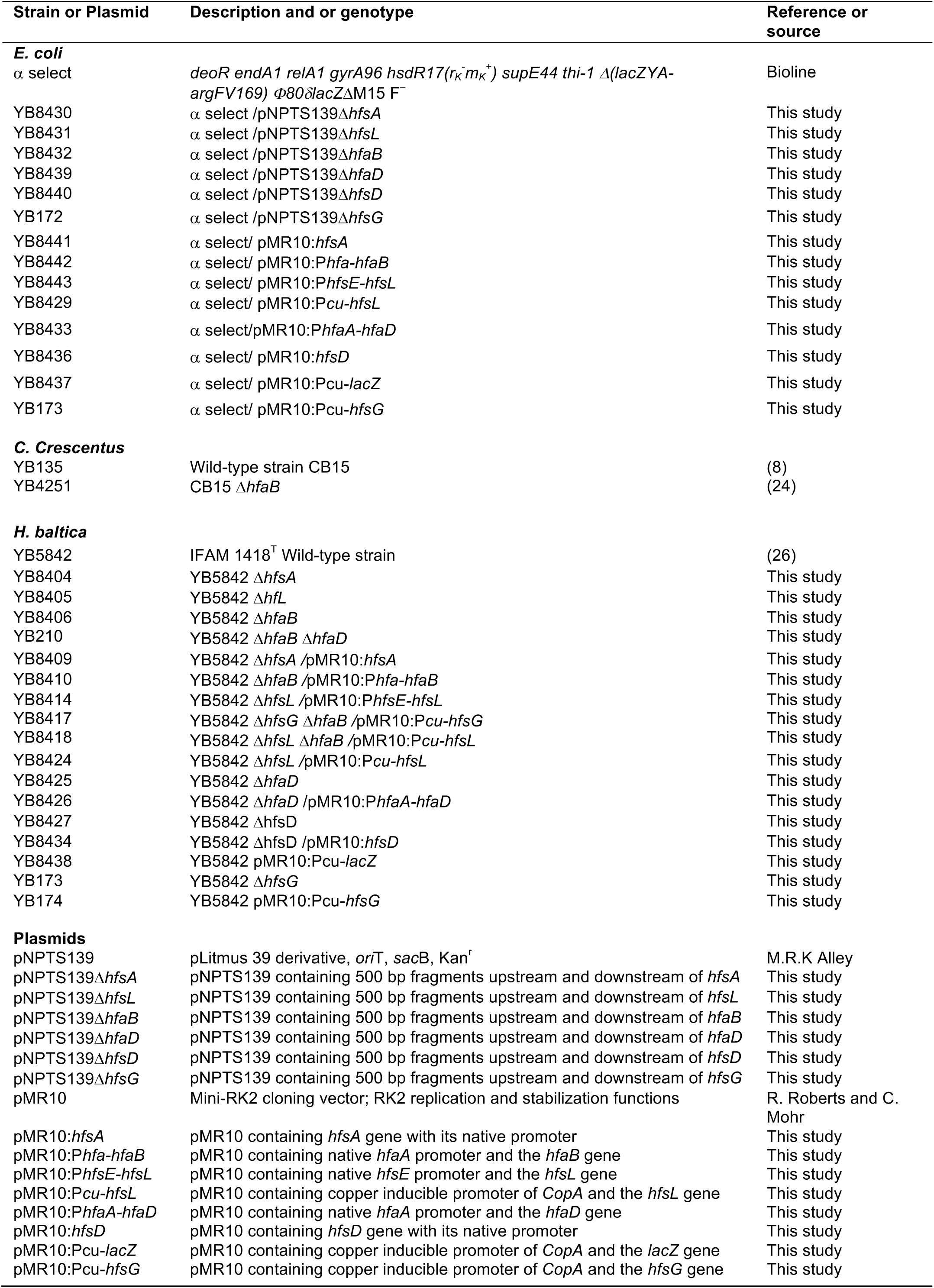
Strains and Plasmids used in this study.

**Table S2:**
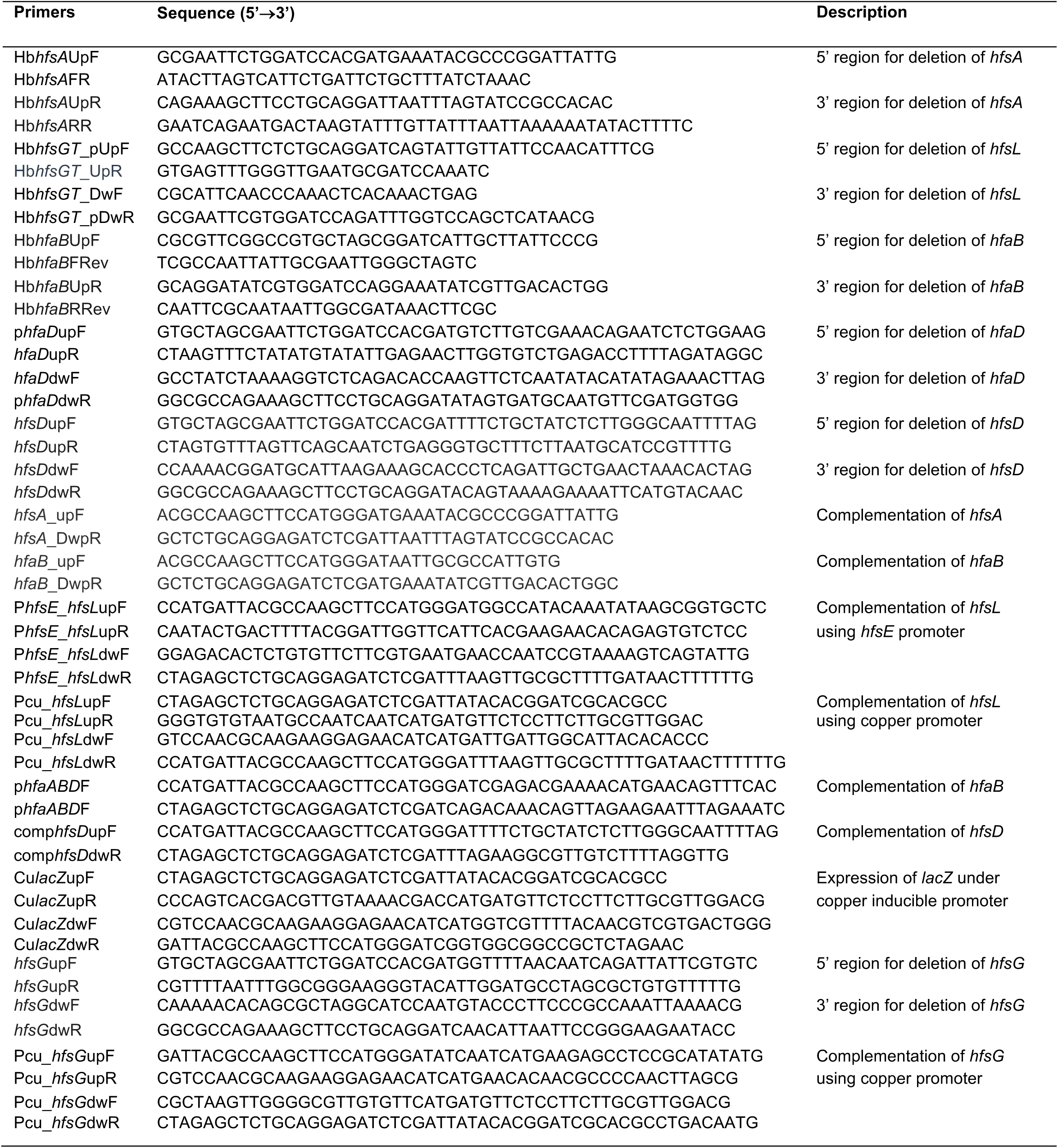
Primers used in this study.

**Table S3:** Lectin binding assays for all the lectins used.

| Lectin | Specificity | *H. baltica* holdfast | *C. crescentus* holdfast |
| --- | --- | --- | --- |
| Wheat Germ Agglutinin | GlcNAc, sialic acid | √ | √ |
| Succinylated Wheat Germ Agglutinin | GlcNAc | √ | √ |
| Lycopersicon Esculentum Tomato | GlcNAc 1-4 | √ | √* |
| Datura Stramonium Lectin | GlcNAc 1-4 | √* | - |
| Solanum Tuberosum Potato Lectin | GlcNAc, prefers trimers and tetramers | √ | √* |
| Ricinus Communis Agglutinin | Galactose | √ | - |
| Griffonia Simplicifolia Lectin 1 | α-GalNAc, α-galactose | √ | - |
| Soybean Agglutinin | α-GalNAc | - | - |
| Concanavalin A | α-linked mannose | - | - |
| Dolichos Biflorus Agglutinin | α-linked acetylgalactosamine | - | - |
| Peanut Agglutinin | Galactosyl β-1,3 N-acetylgalactosamine | - | - |
| Soybean Agglutinin | α or β acetylgalactosamine | - | - |
| Ulex Europaeus Agglutinin 1 | N- acetylgalactosamine, sialic acid or chitobiose | - | - |
| Len Culinaris Agglutinin | α-linked mannose | - | - |
| Pisum Sativum Agglutinin | α-linked mannose, fucose or N-acetylchitobiose | - | - |
| Erythrina Cristagalli Lectin | Galactose, prefers Galactosyl β-1,4 N- acetylgalactosamine | - | - |
| Jacalin | Galactosyl β-1,3 N-acetylgalactosamine | - | - |
| Griffonia Simplicifolia Lectin 2 | α or β acetylgalactosamine | - | - |
| Vicia Villosa Lectin | α or β terminal N-acetylgalactosamine | - | - |

√ Fluorescent signal detected
- No fluorescent signal detected
\* Binding is enhanced on rosettes but weaker signals on single cells.

**Figure S1:**
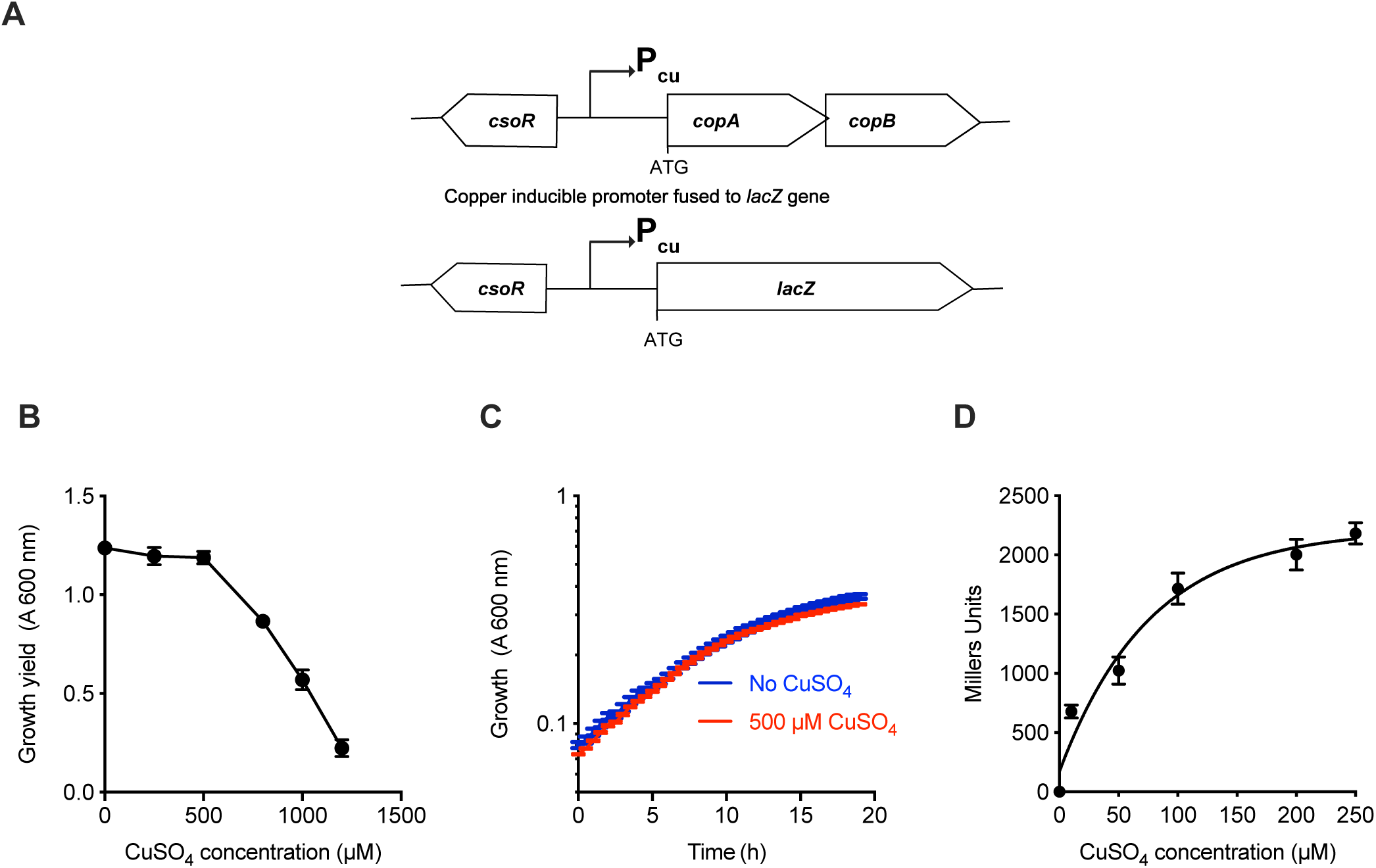
Design of a copper inducible promoter system in *H. baltica*. **A.** Chromosomal arrangement of one of the copper sensitive operon (*Cso*) genes in *H. baltica* genome, showing copper operon repressor *csoR* and copper binding proteins *copA* and *copB* (top panel). The bottom diagram shows the fusion of copper-inducible promoter (P_*cu*_) to *lacZ* reporter gene. **B.** Effect of different concentration of CuSO_4_ added into marine broth on *H. baltica* growth. Growth yield (OD_600_) was measured on overnight cultures with different concentration of CuSO_4_. Data represent mean of four independent replicates and the error bars represent standard error. **C**. Representative growth curves of *H. baltica* growing in marine broth without or with 500 µM CuSO_4_. OD_600_ representing bacterial growth in a 24 well plate was measured every 30 min. **D.** β–galactosidase activity representing the P_cu_ activity when induced with different concentrations of CuSO_4_. Exponential cultures were induced for 4 hrs. Data shown is representative of three independent replicates.

**Figure S2:**
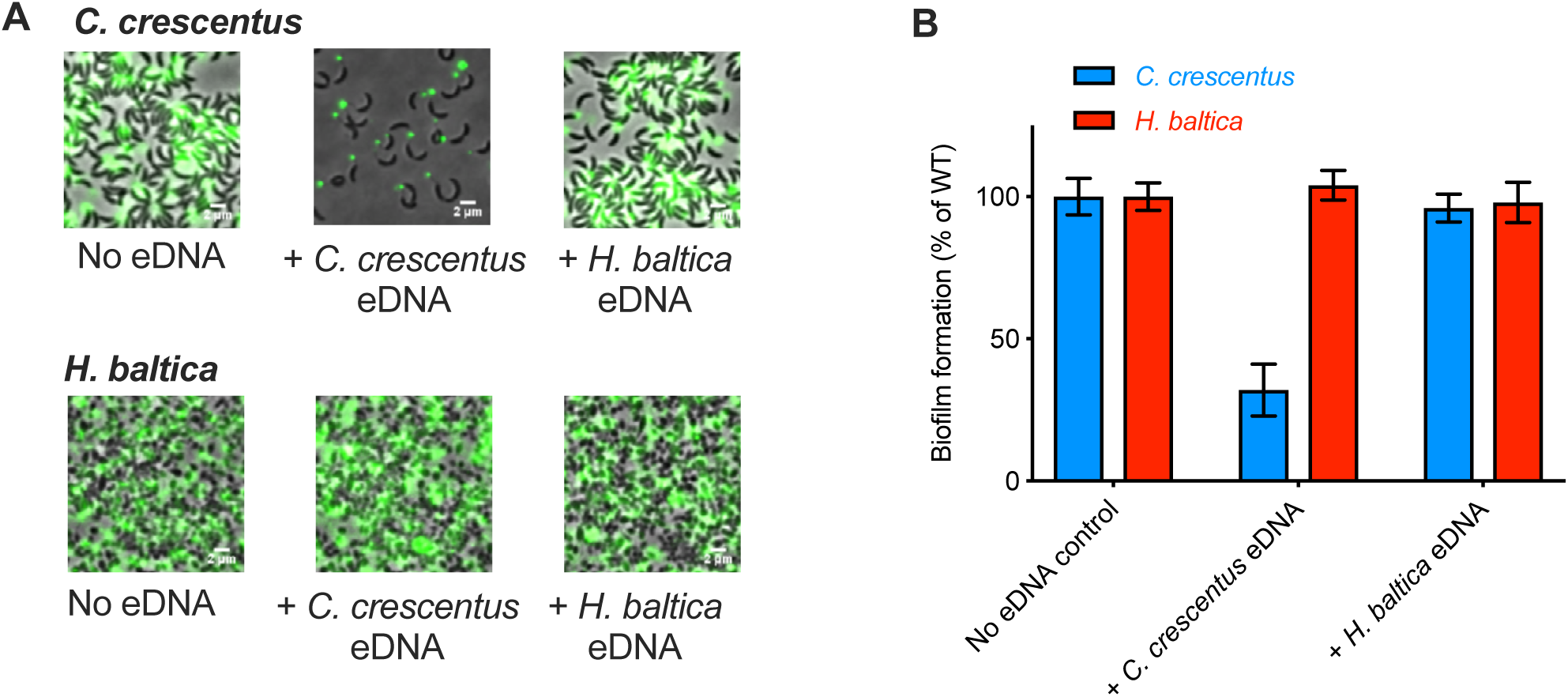
DNA inhibition of holdfast binding and biofilm formation. **A.** *C. crescentus* (upper panel) and *H. baltica* (lower panel) cells bound to a glass surface in presence of eDNA from each strain. Holdfasts labeled with WGA-AF488 lectins after exponentially grown cells were bound to a glass slide for 45 min. **B.** Biofilm quantification after 24 h for *C. crescentus and H. baltica* in presence of eDNA. Data are expressed as an average of 4 independent replicates and the error bars represent the standard error.

